# Scleraxis Lineage Cells Contribute to Organized Bridging Tissue During Tendon Healing, and Identifies Subpopulations of Resident Tendon Cells

**DOI:** 10.1101/469619

**Authors:** Katherine T. Best, Alayna E. Loiselle

**Author notes:** Corresponding Author Alayna E. Loiselle, PhD, Center for Musculoskeletal Research, University of Rochester Medical Center, 601 Elmwood Ave, Box 665, Rochester, NY, 14642.

## Abstract

During tendon healing, it is postulated that intrinsic tendon cells drive tissue regeneration while extrinsic cells drive pathological scar formation. Intrinsic tendon cells are frequently described as a homogenous, fibroblast population that is positive for the marker Scleraxis. It is controversial whether intrinsic Scleraxis localize within the forming scar tissue during adult tendon healing. We have previously demonstrated that calcium binding protein S100a4 is a driver of tendon scar formation and marks a subset of intrinsic tendon cells. However, the relationship between Scleraxis and S100a4 has not been explored. In this study, we aimed to investigate the localization of Scleraxis lineage cells following adult murine flexor tendon repair and to establish the relationship between Scleraxis and S100a4 throughout both homeostasis and healing. We have shown that adult Scleraxis lineage cells localize within the scar tissue and organize into a highly aligned cellular bridge during tendon healing. Additionally, we demonstrate that markers Scleraxis and S100a4 label distinct populations in tendon during homeostasis and localize differently within tendon scar tissue, with Scleraxis found specifically in the organized bridging tissue and S100a4 localized throughout the entire scar region. These studies define a heterogeneous tendon cell environment and demonstrate discreet contributions of subpopulations during healing. Taken together, these data enhance our understanding and ability to target the complex cellular environment of the tendon during homeostasis and healing.

## Introduction

Tendon is a dense connective tissue that transmits force from muscle to bone. The transmission of force is facilitated by the tendon’s structure, which consists of an aligned and organized type I collagen extracellular matrix. Following adult acute tendon injury, tendon function is disrupted by the generation of scar tissue, a highly disorganized extracellular matrix consisting of primarily type III collagen. The scar tissue fails to be fully remodeled during healing, impeding tendon function and increasing the incidence of post-operative tendon rupture. Despite the prevalence of acute tendon injuries, the cellular components and molecular mechanisms driving scar formation during tendon healing have not been extensively characterized.

Localization and function of the various cell populations present during tendon healing are still contested. Better understanding of the intrinsic tendon cell population (i.e. resident tendon fibroblasts) and extrinsic populations (i.e. inflammatory cells, non-tendon mesenchymal cells) during tendon healing could inform identification of biological therapeutics. Fetal and neonatal tendons have increased recruitment of native tendon fibroblasts (tenocytes) to the defect site and exhibit regenerative capacities following an acute injury [1-3], whereas injuries in adult animals exhibit a decreased recruitment of intrinsic cells and heal with scar [3]. Therefore, it is often the extrinsic cells that are implicated in scar tissue formation, increasing interest in how to harness the regenerative capabilities of intrinsic tendon cells. Despite the potential role of tenocytes in tendon regeneration, the localization of these cells during healing is debated.

Tenocytes are typically regarded as a homogenous population positive for the marker Scleraxis (*Scx*). Scx is a basic helix-loop-helix transcription factor that directs expression of extracellular matrix components and is necessary for the proper development of force-transmitting tendons [4-6]. The localization of Scx+ cells during healing differs greatly between specific tendons and injury models, with some studies showing Scx+ cells localized at and bridging the defect site [7, 8] while others exhibit a complete absence of Scx+ cells within the scar [3]. No studies have extensively characterized Scx lineage cell localization in either a flexor tendon injury model or a model of complete transection and repair.

In potential contradiction to the notion of a homogeneous resident tendon cell population, we have recently demonstrated that S100a4 marks a high proportion of intrinsic tendon cells [9, 10]. However, the relationship between the Scx and S100a4 populations is unclear both during tendon homeostasis, and in response to injury. We hypothesized that in a model of complete transection and repair of the flexor tendon, Scx+ populations will localize within the scar tissue and that Scx would mark a separate population from the S100a4 cells during homeostasis and healing. In the current study, we characterized the localization of adult Scx-lineage cells during tendon healing, utilizing different labeling schemes to assess a variety of Scx+ populations. In addition, we have examined the relationship between Scx+ and S100a4+ cells during homeostasis and throughout tendon healing and assessed the potential of Scx-lineage cells to take on a terminal myofibroblast fate.

## Methods

### Animals Ethics

This study was carried out in strict accordance with the recommendations in the Guide for the Care and Use of Laboratory Animals of the National Institutes of Health. All animal procedures were approved by the University Committee on Animal Research (UCAR) at the University of Rochester.

### Mouse Models

Scx-Cre^ERT2^ mice were generously provided by Dr. Ronen Schweitzer. C57Bl/6J (#000664), ROSA-Ai9 (#007909), and S100a4-GFP (#012893) mice were obtained from The Jackson Laboratory (Bar Harbor, ME). ROSA-Ai9 mice express Tomato red fluorescence in the presence of Cre-mediated recombination [11]. S100a4-GFP mice express eGFP under control of the S100a4 promoter [12]. Scx-Cre^ERT2^ mice were crossed to the ROSA-Ai9 to trace Scx-expressing cells (Scx^Ai9^). Scx^Ai9^ animals received three 100mg/kg i.p. tamoxifen injections at the times outlined in all figures.

### Flexor Tendon Repair

At 10-12 weeks of age, mice underwent complete transection and repair of the flexor digitorum longus (FDL) tendon in the hindpaw as previously described [13]. Briefly, mice were injected prior to surgery with 15-20μg of sustained-release buprenorphine. Mice were then anesthetized with Ketamine (60mg/kg) and Xylazine (4mg/kg). Following sterilization of the surgery region, the FDL tendon was transected at the myotendinous junction to reduce strain-induced rupture of the repair site and the skin was closed with a 5-0 suture. A small incision was then made on the posterior surface of the hindpaw, the FDL tendon was located and completely transected using micro spring-scissors. The tendon was repaired using an 8-0 suture and the skin was closed with a 5-0 suture. Following surgery, animals resumed prior cage activity, food intake, and water consumption.

### Histology – Frozen

Hindpaws were harvested from uninjured, 7, 14, and 21 days post-repair for frozen sectioning. Hindpaws were fixed in 10% formalin for 24 hours at 4°C, decalcified in 14% EDTA for 4 days at 4°C, and processed in 30% sucrose for 24 hours at 4°C to cryo-protect the tissue. Samples were then embedded in Cryomatrix (ThermoFisher, Waltham, MA) and sectioned into 8µm sagittal sections using a cryotape-transfer method [14]. Sections were mounted on glass slides using 1% chitosan in 0.25% acetic acid and counterstained with the nuclear stain DAPI. Endogenous fluorescence was imaged on a VS120 Virtual Slide Microscope (Olympus, Waltham, MA). Images are representative of 3-4 specimens per time-point.

### Histology – Paraffin

Hindpaws were harvested from C57Bl/6J mice for uninjured, 7, 14, 21, and 28 days post-repair time-points and from Scx^Lin^ mice at day 14 post-repair for paraffin sectioning. Hindpaws were fixed in 10% formalin for 72 hours at room temperature, decalcified in 14% EDTA for two weeks, processed and embedded in paraffin. Three-micron sagittal sections were cut, de-waxed, rehydrated, and probed with antibodies for Scx (1:100, Cat#: ab58655, Abcam, Cambridge, MA), Red fluorescent protein (RFP) (1:500, Cat#: ab62341, Abcam, Cambridge, MA) and αSMA-CY3 (1:200, Cat#: C6198, Sigma Life Sciences, St. Louis, MO). A Rhodamine Red-X AffiniPure secondary antibody was used to fluorescently label RFP (1:200, Cat#: 711-296-152, Jackson ImmunoResearch, West Grove, PA). Nuclei were counterstained with DAPI and imaging was performed using a VS120 Virtual Slide Microscope (Olympus, Waltham, MA).

## Results

### Scleraxis lineage cells contribute to highly organized bridging tissue within scar during flexor tendon repair

To label the Scleraxis lineage cell population prior to injury, Scx^Ai9^ mice were injected with tamoxifen for three days followed by a four day washout period immediately prior to surgery (Scx^Lin^). Specimens were harvested at days 7, 14, and 21 post-surgery (Fig. 1A & B). In uninjured tendons, Scx^Lin^ cells represented the majority of the total tendon cell population (Fig. 1C). By 7 days following repair, Scx^Lin^ cells resided within the tendon body and were not found in the scar tissue. The epitenon expanded in size and remained absent of Scx^Lin^ cells (Fig. 1D). By day 14, Scx^Lin^ cells were organized into an aligned bridging tissue that spanned the width of the scar tissue and these cells appeared to emanate from the epitenon, which also comprised of Scx^Lin^ cells (Fig. 1E & E’). At 21 days post-repair, Scx^Lin^ cells were still localized to the tendon stubs and bridging scar tissue and had acquired a wavy morphology (Fig. 1F & F’).

**Figure 1.**
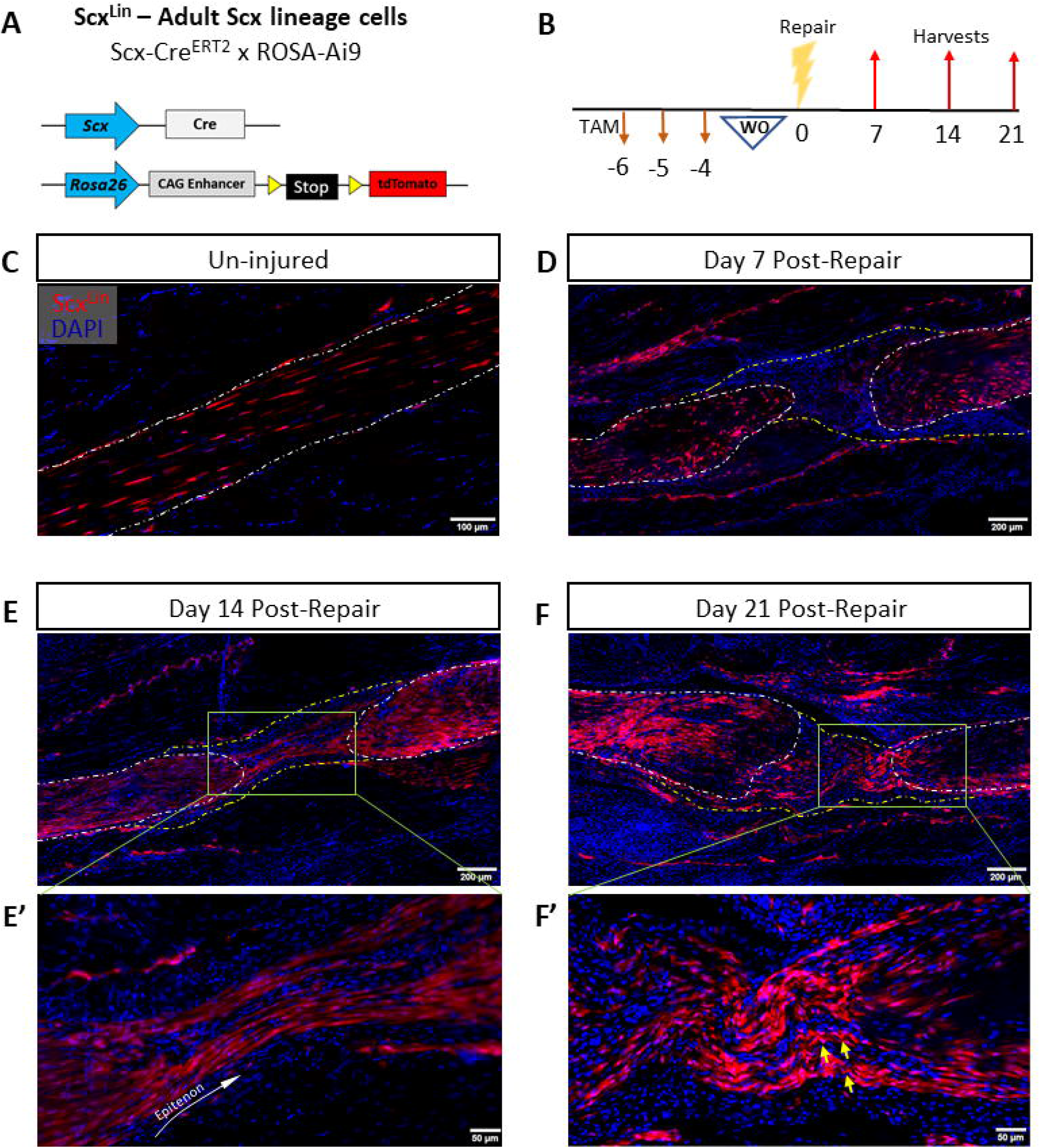
Scleraxis lineage cells localize within the forming scar tissue during flexor tendon healing. Scx-Cre^ERT2^ mice were crossed to ROSA-Ai9 reporter (Scx^Ai9^) to trace adult Scx lineage cells at homeostasis and throughout healing (Scx^Lin^) (A). Mice were injected with tamoxifen (TAM) for three consecutive days and tamoxifen was allowed to washout (WO) for four days prior to tendon injury and repair (B). Tendons were harvested uninjured (C) and at days 7 (D), 14 (E & E’), and 21 (F & F’) post-repair. Nuclear stain DAPI is blue. Tendon is outlined by white dotted lines and scar tissue by yellow dotted lines. White arrows indicate Scx^Lin^ with wavy morphology.

### Scleraxis Expression is Biphasic During Early Tendon Healing

To assess the localization of cells expressing Scleraxis immediately after injury, mice were injected with tamoxifen on days 0-2 following repair (Fig. 2A). The localization of Scx^Ai9^ cells in the uninjured contralateral control tendon when labelling was performed from days 0-2 post-surgery (Scx^0-2^) was comparable to Scx^Ai9^ expression when labelling was performed prior to surgery (Scx^lin^) (Fig. 2B). No Scx^0-2^ cells were found within the scar tissue in three out of four samples and there were fewer Scx^0-2^ cells altogether compared to Scx^Lin^ (Fig. 2C & C’). To assess localization of cells expressing Scleraxis in the early proliferative stage of healing, mice were injected with tamoxifen on days 5-7 following repair (Fig. 2D). Localization of Scx^Ai9^ cells in the contralateral control with Scx^5-7^ tendon was also comparable to Scx^Lin^ (Fig. 2E). At 14 days following repair, Scx^5-7^ cells were found within the scar tissue at the injury site, similar to Scx^Lin^ (Fig. 2F & F’). Thus, different induction schemes label different tendon cell populations, suggesting changes in Scx expression post-repair.

**Figure 2.**
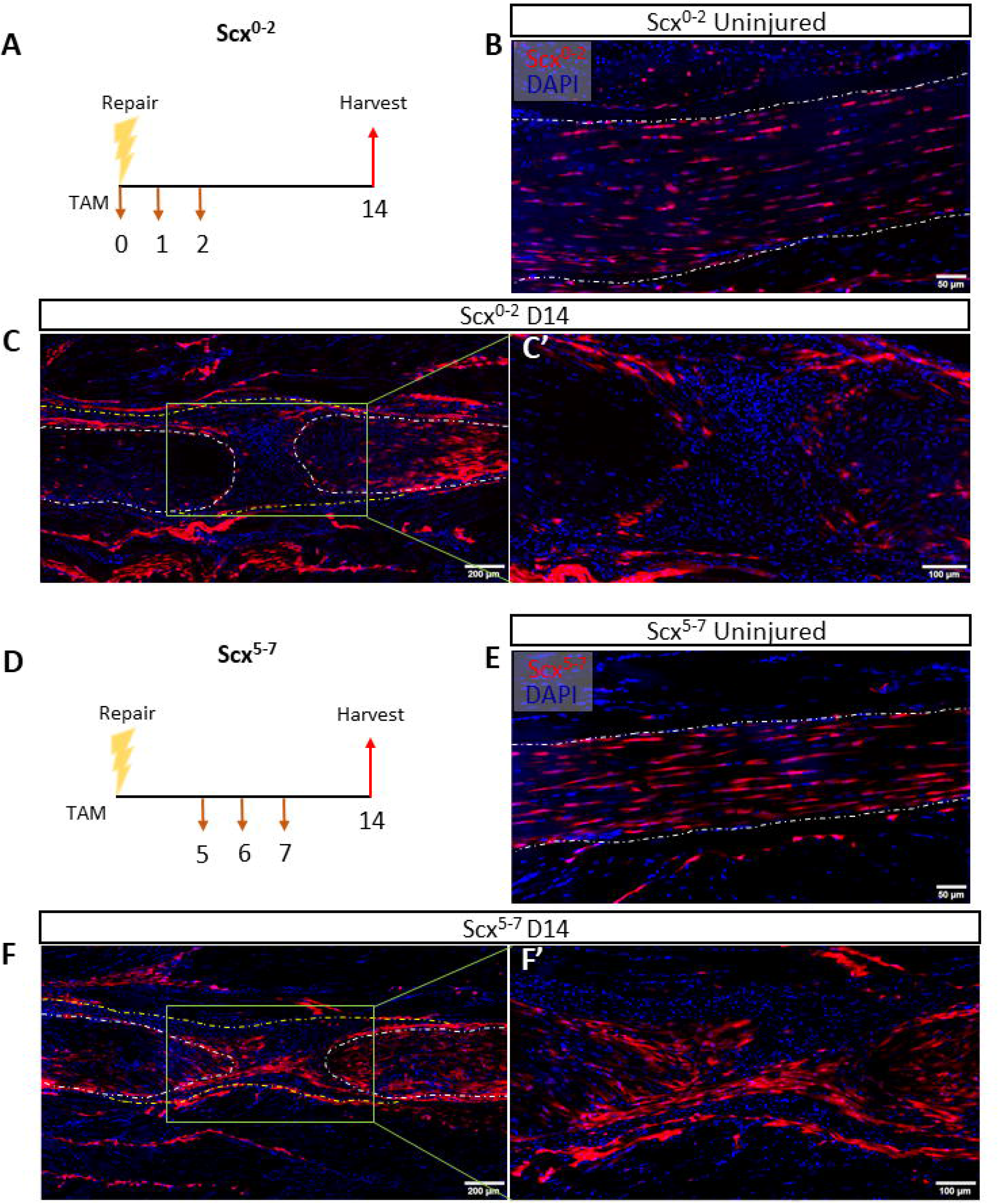
Scleraxis expression is biphasic in early tendon healing. Scx^Ai9^ mice were injected with TAM on days 0-2 post-repair to trace cells expressing Scx shortly after repair (Scx^0-2^) (A). Scx^0-2^ mice were harvested uninjured (B) and 14 days post-repair (C & C’). Scx^Ai9^ mice were injected with TAM on days 5-7 post-repair to trace cells expressing Scx later in healing (Scx^5-7^) (D). Scx^5-7^ mice were harvested uninjured (E) and 14 days post-repair (F & F’). Nuclear stain DAPI is blue. Tendon is outlined by white dotted lines and scar tissue by yellow dotted lines.

### Scleraxis and S100a4 label distinct and overlapping tendon cells, demonstrating that tenocytes are a heterogenous population

Scx^Lin^ cells clearly mark an adult resident tendon cell population, and we have previously demonstrated that S100a4 is also expressed by many of these cells [9, 10], however, the relationship between these two populations is unclear. To address this, Scx^Ai9^ mice were crossed to S100a4^GFP^ mice to generate Scx^Ai9^S100a4^GFP^ animals (Fig 3A). In this model, cells expressing Scx fluoresce red following tamoxifen injection while all cells actively expressing S100a4 fluoresce green. To assess the resident tendon cell populations independent of injury, uninjured tendons were used. Mice were injected with tamoxifen three times and allowed to washout for four days before harvest (Scx^Lin^S100a4^GFP^) (Fig 3B). Four individual tendon cell populations were identified: Scx^Lin^, S100a4^GFP^, Scx^Lin^;S100a4^GFP^, and a dual-negative population. This data highlights previously unappreciated heterogeneity within the adult tendon cell population.

**Figure 3.**
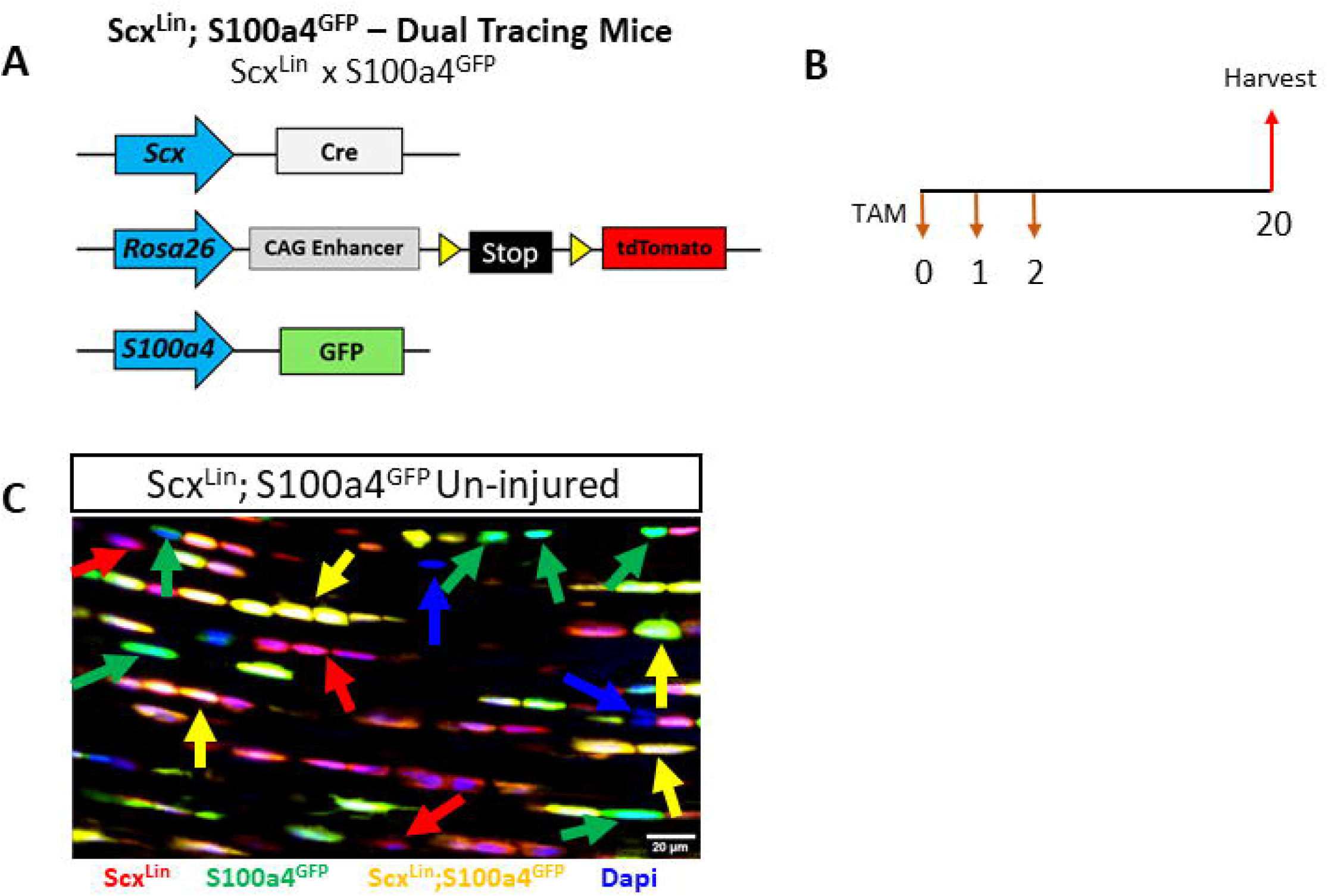
Dual tracing of Scx^Lin^ and S100a4^GFP^ cells exhibits tendon cell heterogeneity. Scx^Ai9^ mice were crossed to S100a4^GFP^ mice (Scx^Ai9^S100a4^GFP^) to allow dual tracing of Scleraxis lineage and S100a4+ cell populations (A). Scx^Ai9^S100a4^GFP^ mice were injected with TAM on three consecutive days and harvested 18 days following the final injection (B). Uninjured Scx^Ai9^S100a4^GFP^ tendons exhibit four distinct cell types with colored arrows indicating examples of each: Scx^Lin^ (red arrow), S100a4^GFP^ (green arrow), Scx^Lin^; S100a4^GFP^ (yellow arrow), and dual-negative populations (blue arrow) (C). Nuclear stain DAPI is blue.

### Scleraxis and S100a4 cell populations localize differently during flexor tendon repair

We next sought to characterize the localization of these cell populations relative to one another during flexor tendon healing. Mice were injected with tamoxifen using the washout scheme (Fig 4A). At 14 days post-repair, Scx^Lin^ cells were primarily localized to the tendon stubs and periphery of the scar tissue while S100a4^GFP^ cells were localized throughout the entire scar tissue area. Scx^Lin^;S100a4^GFP^ cells were present at the tendon stubs and sparsely in the scar tissue. (Fig 4B). By 21 days post-repair, the Scx^Lin^ population had expanded and was specific to the bridging scar tissue and tendon stubs. Interestingly, S100a4^GFP^ cells were found both within the organized bridging scar tissue exhibiting an elongated morphology and throughout the disorganized scar tissue with a rounded morphology. Dual-expressing Scx^Lin^;S100a4^GFP^ cells were found prevalently in the bridging tissue at 21 days post-repair (Fig 4C, C’ & C”). To assess if some of these dual-expressing cells were still actively expressing Scx throughout healing, we injected Scx^Ai9^;S100a4^GFP^ mice with tamoxifen at days 10-12 to label cells expressing Scx nearly two weeks into healing (Scx^10-12^) (Fig 4D). When harvested at 14 days post-repair, Scx^10-12^ cells were still present within the tendon stubs and bridging tissue. Some of these cells were positive for both markers, indicating that a subset of cells later in healing are still expressing both Scx and S100a4 (Fig 4E).

**Figure 4.**
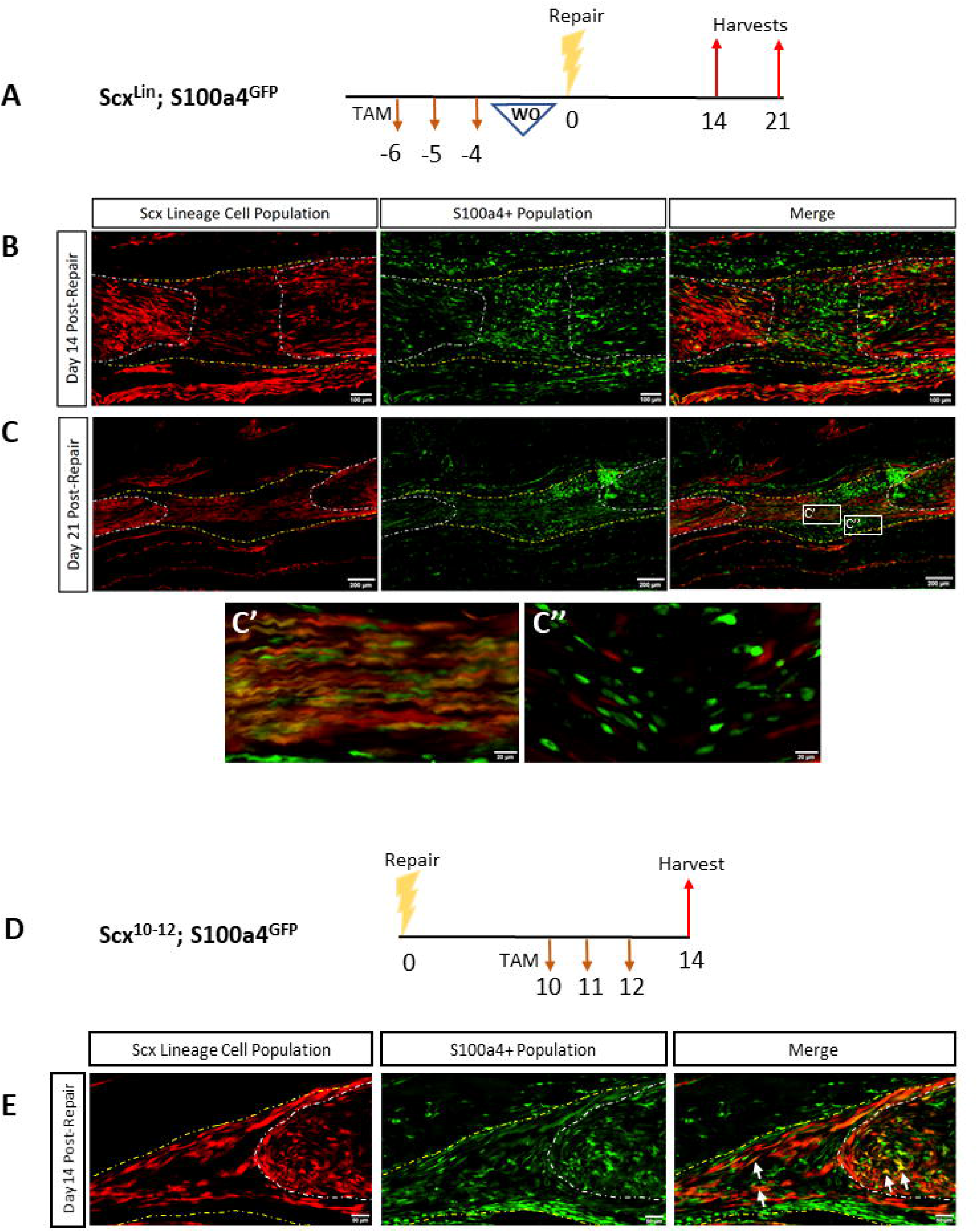
Scx^Lin^ and S100a4^GFP^ populations differentially localize during tendon healing. Scx^Ai9^; S100a4^GFP^ mice were injected with TAM on three consecutive days and allowed to washout for four days prior to repair (A). Scx^Lin^; S100a4^GFP^ mice were harvested at 14 (B) and 21 (C) days post-repair. Insets indicate linear (C’) and rounded (C”) morphology of S100a4^GFP^ cells. Scx^Ai9^; S100a4^GFP^ mice were injected with TAM on days 10-12 post-repair (D) to generate Scx^10-12^; S100a4^GFP^ mice and were harvested at day 14 (E). Tendon is outlined by white dotted lines and scar tissue by yellow dotted lines.

### Scleraxis lineage cells do not primarily differentiate into myofibroblasts during healing

It has previously been shown that Scx can drive αSMA+ myofibroblast differentiation in the heart [15]. To investigate the ability of Scx^Lin^ cells in the tendon to differentiate into myofibroblasts, we performed co-immunofluorescence of RFP and αSMA on day 14 post-repair sections (Fig 5A). A small proportion of Scx^Lin^ cells expressed αSMA within the tendon stub, with nearly no Scx^Lin^ cells expressing αSMA within the bridging scar tissue (Fig 5B). Additionally, most αSMA+ myofibroblasts were not expressing RFP, suggesting that the majority of myofibroblasts are not derived from the Scx^Lin^ cells.

**Figure 5.**
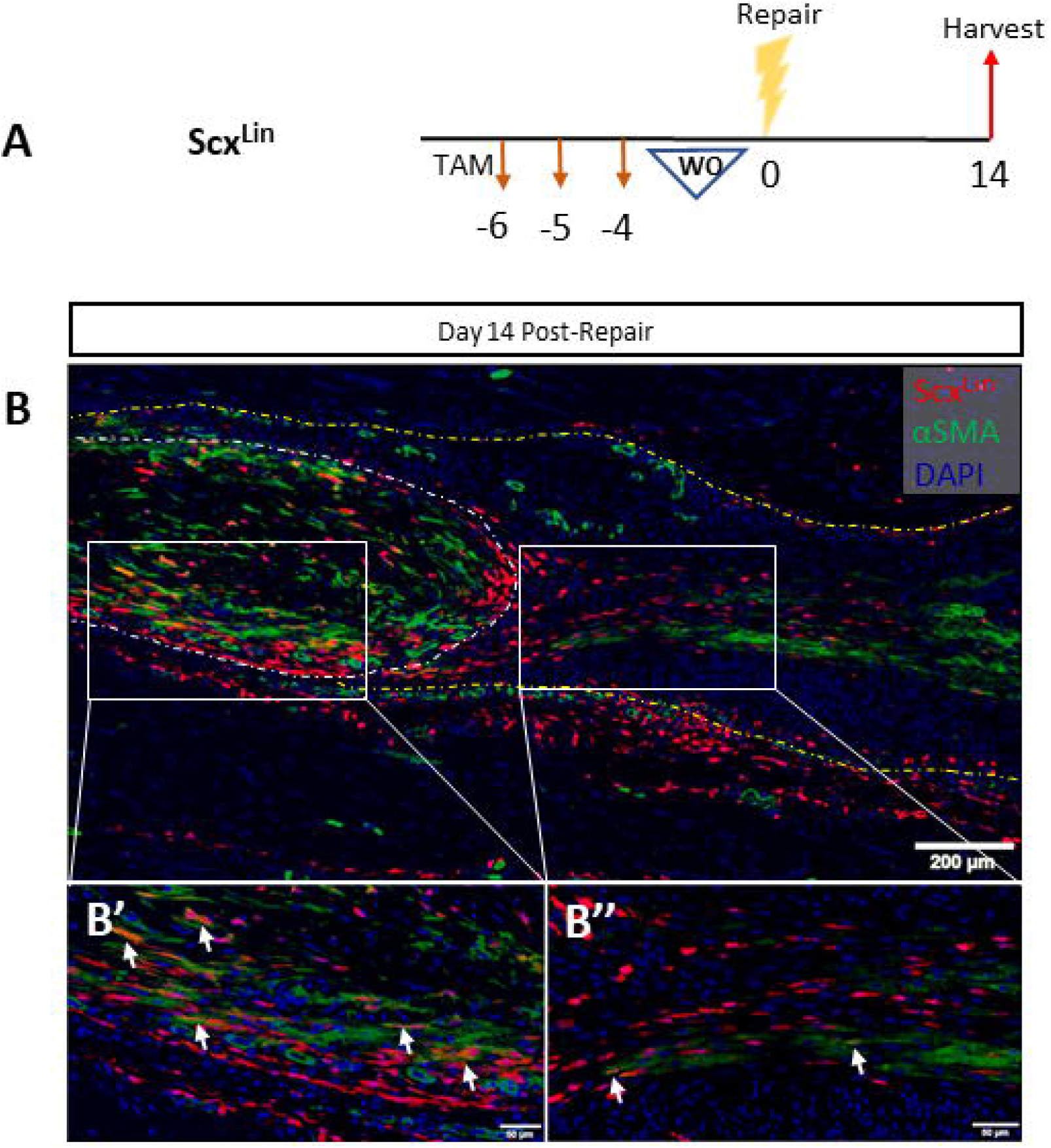
Scx^Lin^ do not substantially differentiate into myofibroblasts. Scx^Ai9^ mice were injected with tamoxifen for three days and allowed to washout for four days (Scx^Lin^) (A). Coimmunofluorescence of red fluorescent protein (RFP, labels Scx^Lin^ cells) and αSMA (myofibroblast marker) on Scx^Lin^ day 14 post-repair sections reveals minimal differentiation of Scx^Lin^ cells into myofibroblasts (B). Differentiation occurs primarily within the native tendon (B’) and not within the scar tissue (B”). Nuclear stain DAPI is blue. Tendon is outlined by white dotted lines and scar tissue by yellow dotted lines.

## Discussion

In the present study, we assessed the localization of Scx lineage cells throughout flexor tendon healing and examined the relationship between Scx and S100a4 cell populations during tendon homeostasis and healing. We have established that Scx lineage cells localize within the scar tissue in a highly aligned and organized manner with Scx expression occurring in a biphasic pattern, in a model of complete transection and repair of the flexor tendon. We have also shown that while Scx and S100a4 are both expressed in a subset of resident tendon cells, there are also populations that express only one of these markers, as well as a double negative population, thus identifying much greater heterogeneity of the resident tendon cell population than previously appreciated. In addition, Scx+ and S100a4+ cells localize differently during tendon injury, suggesting differential roles for these cells during tendon healing.

Previous studies focused on elucidating the localization of Scx cell populations during tendon healing have yielded conflicting findings. In a patellar tendon window-defect model, it was demonstrated that cells from the surrounding paratenon proliferated, migrated to the defect site, and became Scx+ by 7 days post-injury, with Scx+ cells bridging the injury site by 14 days [7]. Additionally, it has been demonstrated that a subset of αSMA+ lineage cells derived from the paratenon bridge the patellar tendon window injury and become Scx+ 14 days following repair [16]. In contrast, a complete transection injury model of the Achilles tendon demonstrated that while Scx+ cells were present in the gap space and could induce regeneration of neonatal tendon, there was a nearly complete absence of Scx+ and Scx lineage cells in the scar tissue formed within the adult tendon gap space [3]. Interestingly, a recent study by Sakabe *et al*. demonstrated that a partial Achilles tendon transection had Scx+ cells within the defect site by day 14 post-injury [8]. This suggests that keeping the cut ends of the tendon near one another and the maintenance of tensile force across the defect may be necessary for localization of Scx+ cells within the defect. Consistent with this, while our repair model does result in complete transection of the tendon, the surgical repair both maintains tensile force and tendon stub proximity, likely facilitating the migration of Scx-lineage cells in to the bridging tissue of the defect. It has previously been shown that tendon cells lose Scx expression shortly after acute injury ([8]) and that Scx expression is mechanosensitive [17]. Traces of Scx^0-2^ and Scx^5-7^ populations revealed biphasic Scx expression patterns during healing, likely resulting from the initial loss of physiological tension at the injury site.

It has previously been shown that epitenon fibroblasts become highly Scx+ and migrate from the epitenon into the tendon fascicles following addition of physiological loading via treadmill running [18]. In addition, Dyment et al., demonstrate that Scx+ cells originating from the paratenon contribute to tissue bridging in a patellar tendon defect model (dyment papers). Consistent with this, we observed that the outer portion of the flexor tendon, known as the epitenon, had expanded in size and was positive for Scx^Lin^ cells, with these Scx^Lin^ contributing to the bridging tissue. Identification of epitenon specific markers will permit better understanding of the function and spatial contribution of epitenon-derived cells.

A trace of adult Scx lineage cells revealed that not all tenocytes are Scx^Lin^ during homeostasis, providing evidence that tenocytes are not a homogenous population. It has previously been shown that Scx-GFP mice [19] label most, but not all, tendon cells [8]. Additionally, previous work has shown that αSMA lineage cells mark a subset of Scx+ tendon cells in uninjured patellar tendon [16]. We have previously shown that the calcium binding protein S100a4, which has been implicated in pathological fibrosis in a variety of tissues, labels a subset of intrinsic tendon cells and is present within the scar tissue during flexor tendon healing [9, 10]. Additionally, conditional deletion of *EP4* in S100a4-lineage and Scx-lineage cells results in distinct phenotypes during tendon healing, suggesting these may represent discreet cell populations [9]. Consistent with this, dual tracing of Scx and S100a4 demonstrated that there are at least four distinct tendon cell sub-populations during homeostasis: Scx^Lin^, S100a4^GFP^, Scx^Lin^;S100a4^GFP^, and a dual-negative population. Taken together, these findings demonstrate that intrinsic tendon cells are not a homogenous population. While future studies will be needed to delineate the potential discreet functions of these populations during tendon homeostasis, the dual-negative cell population may represent a non-fibroblast resident cell population.

Our data also suggest some divergence in the spatial localization of S100a4 and Scx cells during healing, suggesting they may be performing divergent functions. By 21 days post-repair, the Scx^Lin^ cells were highly specific to the tendon stubs and bridging region of the scar tissue. Interestingly, the S100a4 cell population is found both within the organized bridging scar tissue with an elongated cellular morphology similar to Scx^Lin^ cells, and within the peripheral scar tissue exhibiting a rounded morphology. The relative contribution of tendon-derived S100a4^+^ cells and extrinsic S100a4^+^ cells is currently unknown, however, the differences in cellular morphology may be indicative of discreet S100a4 populations that originate from different sources and perform divergent functions during healing.

Scx drives expression of extracellular matrix components, including fibrillar collagens and fibronectin [4, 5, 20]. In pathological settings, fibroblasts can differentiate into contractile myofibroblasts that produce extracellular matrix proteins and aid in wound closure [21, 22]. It has been shown that Scx expression in cardiac fibroblasts directly transactivates the α-SMA promoter, resulting in α-SMA+ myofibroblast differentiation [15]. Recently, it has been demonstrated that Scx can also directly transactivate the Twist1 and Snai1 promoters, two transcriptional regulators of epithelial mesenchymal transition, further suggesting that Scx can drive myofibroblast differentiation in cardiac fibroblasts [23]. However, the effects of Scx-induced αSMA expression are tissue specific as it has previously been shown that Scx represses αSMA expression in the mesangial cells of the kidney. In our flexor tendon repair model, we found that a subset of Scx lineage cells differentiated into myofibroblasts [24]. However, the majority of myofibroblasts were derived from a different source, suggesting that Scx may drive the differentiation of another cell population into myofibroblasts. We have recently shown that a subset of S100a4 lineage (S100a4^Lin^) cells differentiate into αSMA+ myofibroblasts following tendon injury and repair [10]. However, S100a4^GFP^ cells are not αSMA+, suggesting that S100a4^Lin^ cells may be downregulating S100a4 and differentiating into myofibroblasts following injury [10]. The influence of Scx on S100a4 myofibroblast differentiation has not been evaluated.

This study clearly demonstrates that Scx^Lin^ cells localize within the scar tissue following flexor tendon repair and that Scx^Lin^ cells represent a separate population from the S100a4^GFP^ cells, highlighting tendon cell heterogeneity. However, there are a few limitations that must be considered. First, while we administered several tamoxifen induction schemes to capture varying Scx+ cell populations, a complete assessment of current Scx expression was not evaluated. Secondly, we did not look any later than day 21 post-repair, and therefore did not investigate the localization of Scx^Lin^ and S100a4^GFP^ later in healing. Lastly, for our Scx^Lin^ trace we allow four days for tamoxifen washout before tendon repair. The half-life of tamoxifen is approximately 12-16 hours [25, 26], resulting in 0.39-1.56% of the original tamoxifen still residing in the Scx^Lin^ mice at the time of tendon repair. As a result of these trace amounts of tamoxifen, we cannot state with complete certainty that the Scx^Lin^ trace is not labeling a few newly activated Scx+ cells. However, when tamoxifen was administered and allowed to washout for four weeks (Scx^Lin,4W^), the Scx^Lin,4W^ population was comparable to the Scx^Lin^ data, suggesting that 4 days is sufficient for tamoxifen washout (Fig S1). Additionally, Scx^Lin^ and Scx^0-2^ exhibit different localization patterns, further suggesting that a short tamoxifen washout period is adequate.

Altogether, these data demonstrate that the Scx^Lin^ cell population localize within the scar tissue and form a highly aligned bridging tissue that resembles native tendon in its organization. This contrasts with the S100a4^GFP^ population that is disorganized and present throughout the entire scar tissue. This suggests that the intrinsic Scx^Lin^ cells may be driving organized regenerative healing while S100a4^GFP^ cells contribute to the formation of scar. Designing strategies to harness intrinsic cells during adult tendon repair may help in driving tendon regeneration, and a better understanding of the discrete intrinsic cell populations within tendon will help inform on targeted therapies to positively affect healing.

## Figure Legends

**Supplemental Figure 1.**
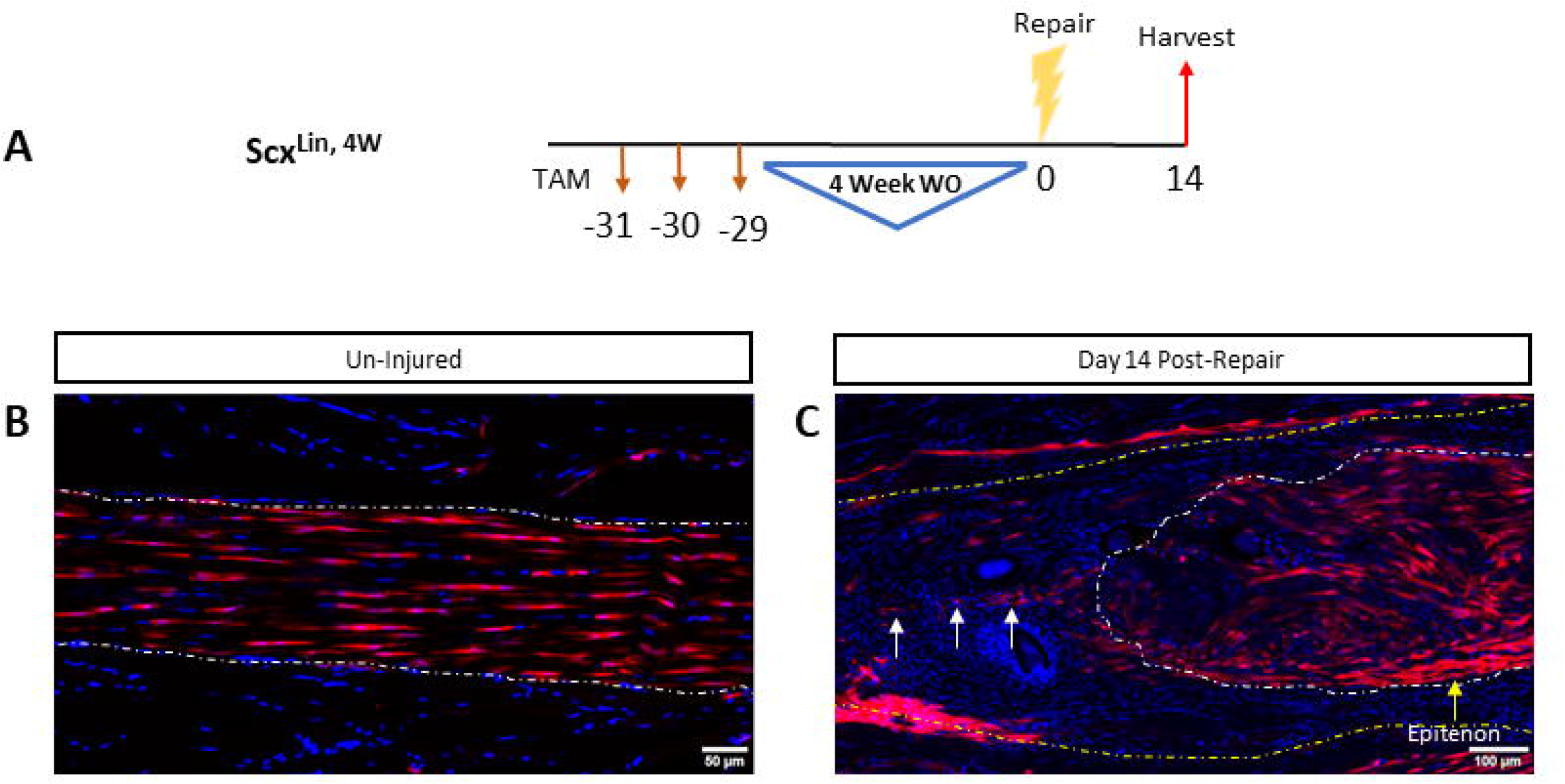
One month tamoxifen washout is comparable to a four day washout. Scx^Ai9^ mice were injected with tamoxifen for three days and allowed to washout for four weeks (Scx^Lin,4W^) (A). The uninjured (B) and 14 day post-repair (C) sections were comparable to Scx^Lin^ mice. White arrows indicate bridging Scx^Lin,4W^ cells and the yellow arrow indicates the enlarged, Scx+ epitenon. Nuclear stain DAPI is blue. Tendon is outlined by white dotted lines and scar tissue by yellow dotted lines.

